# Experimental characterization of *in silico* red-shift predicted iLOV^L470T/Q489K^ and iLOV^V392K/F410V/A426S^ mutants

**DOI:** 10.1101/2021.03.25.436974

**Authors:** Pierre Wehler, Daniel Armbruster, Andreas Günter, Erik Schleicher, Barbara Di Ventura, Mehmet Ali Öztürk

## Abstract

iLOV is a flavin mononucleotide-binding fluorescent protein used for *in vivo* cellular imaging similar to the green fluorescent protein. To expand the range of applications of iLOV, spectrally tuned red-shifted variants are desirable to have reduced phototoxicity and better tissue penetration. In this report, we experimentally tested two iLOV mutants, iLOV^L470T/Q489K^ and iLOV^V392K/F410V/A426S^, which were previously computationally proposed by Khrenova et al. (DOI: 10.1021/acs.jpcb.7b07533) to have red-shifted excitation and emission spectra. We found that mutants bearing the V392K mutation lost the ability to bind FMN. While iLOV^L470T/Q489K^ is about 20% brighter compared to WT iLOV *in vitro*, it exhibits a blue shift in contrast to QM/MM predictions. Additionally, both mutants are expressed at low levels and have undetectable fluorescence in living cells, which prevents their utilization in imaging applications. Our results suggest that rational fluorescent protein design efforts can benefit from *in silico* protein stability and ligand affinity calculations.

## Introduction

Fluorescent proteins (FPs) have revolutionised cell biology by enabling researchers to investigate dynamic cellular processes in real time^1^. The most widely used FP is the green fluorescent protein (GFP) and its spectrally shifted and/or engineered variants^1^, for instance further optimised to more rapidly mature (sfGFP)^2^ or more brightly fluoresce (mNeonGreen)^3^. Despite their usefulness, FPs of the GFP family have some limitations: they depend on molecular oxygen for chromophore formation, which impedes their use to visualise processes under hypoxic or anoxic conditions^4–6^; are relatively large (~25 kDa), which can be problematic in some applications^7,8^, and are mostly sensitive to pH9. While pH-resistant GFP variants have been engineered^9^, oxygen-dependence and size are mostly unchangeable features. FMN-binding fluorescent proteins (FbFPs) overcome these limitations and thus are valuable alternatives to GFP and its variants^10,11^. FbFPs consist of a small light oxygen voltage (LOV) domain (~11 kDa) modified to emit green fluorescence when light in the UV-A-blue range reaches the non-covalently bound FMN chromophore^12^.

One prominent FbFP is iLOV, a derivative of the LOV2 domain of *Arabidopsis thaliana*^7^. A mutation in a key cysteine residue and several rounds of DNA shuffling led to fluorescence emission^7^. While FbFPs have been used in molecular imaging for more than a decade^13^, they suffer from the autofluorescence signal of flavin molecules in the cell and relatively weak fluorescent intensity, issues which have not been improved until very recently^14,15^. Engineering a red-shifted iLOV would bring several advantages, for instance lower phototoxicity and deeper tissue penetration. Moreover, a red-shifted FbFP variant could be used orthogonally to other FbFPs and would potentially allow for multicolor imaging and FRET-based biosensors.

Based on the observation that iLOV possesses several positively charged residues in close proximity to the chromophore, which is a recurrent motif of red fluorescent proteins such as Rtms5^16^ and mKeima^17^, Khrenova and colleagues previously applied quantum mechanics/molecular mechanics (QM/MM) simulations and proposed that the iLOV^Q489K^ mutant would have a ~50 nm red shift in its excitation and emission maxima compared to wild-type (WT) iLOV^18^. Their rationale was that, introducing a positively charged amino group at position 489 next to the chromophore would stabilise the π-electron system of the FMN in the excited state. However, later Davari and colleagues computationally and experimentally showed that K489 is mostly populated in an open conformer, which is far from the chromophore and in fact iLOV^Q489K^ is blue-shifted^19^. In a follow up study, Khrenova and colleagues applied a second round of QM/MM calculations and found other mutations that were predicted to have more stabilised lysine residues next to the chromophore compared to iLOV^Q489K^, which led them to propose iLOV^L470T/Q489K^ and iLOV^V392K/F410V/A426S^ as mutants with ~50 nm red shift for both excitation and emission spectra^20^ (Figure 1). A shift of this extent would open up new potential applications for iLOV and FbFPs in general. In this short report, we experimentally tested these two mutants by measuring the absorption, excitation and emission spectra and calculating the quantum yield and brightness of the purified proteins.

**Figure 1:**
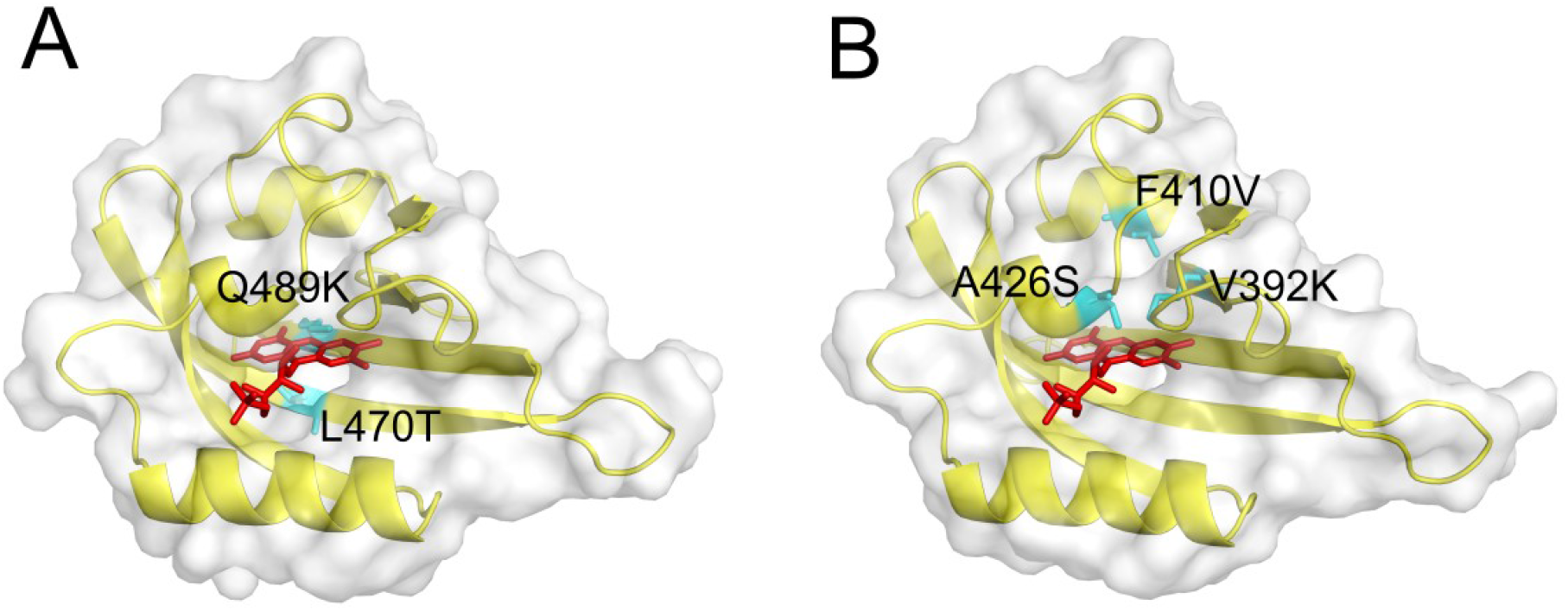
Model structures of iLOV^L470T/Q489K^ and iLOV^V392K/F410V/A426S^ mutants. Cartoon and surface representations of the iLOV^L470T/Q489K^ (A) and iLOV^V392K/F410V/A426S^ (B) mutants. Wild type residues are shown in cyan. The structures (yellow) were generated by the Swiss-Model web-server^21^ and the FMN structures (red) were placed by aligning model structures to the WT iLOV X-ray structure (PDB id: 4EEP). Figures were generated using PyMOL^22^.

## Results

### iLOV^V392K^, iLOV^L470T/Q489K^, and iLOV^V392K/F410V/A426S^ are not detectable in living cells

We started by expressing iLOV^V392K^, iLOV^L470T/Q489K^ and iLOV^V392K/F410V/A426S^ in *E. coli* cells to assess their fluorescence. We used GFP and WT iLOV as positive controls. We found that none of the mutants was detectable under the microscope, while the controls were (Figure 2). We then expressed the same iLOV constructs in mammalian cells, but also here no fluorescence was observed except for WT iLOV (Figure S1).

**Figure 2:**
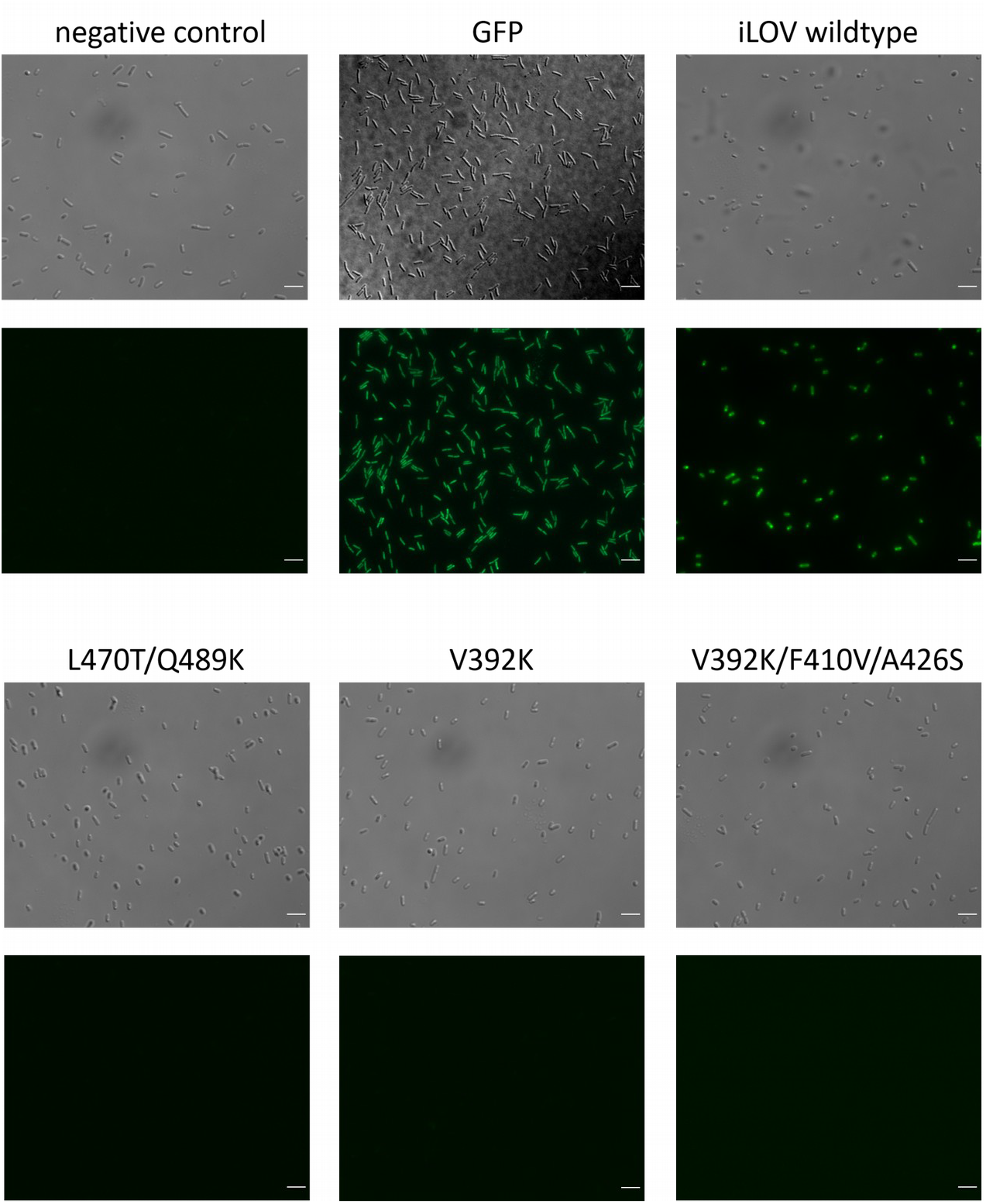
Analysis of the fluorescence of iLOV mutants in bacteria. Representative bright field (top) and GFP channel (bottom) images of *E. coli* Rosetta(DE3) pLysS cells transfected with the indicated constructs. Scale bar for all micrographs is 5 μm. Nontransformed cells were used as negative control. Cells transformed with a plasmid expressing GFP and WT iLOV, respectively, served as positive controls.

One possibility for the lack of fluorescence could be lack of expression. Alternatively, the proteins might be well expressed, but might have lost the ability to bind the chromophore FMN. Similarly, only GFP and WT iLOV cell pellets exhibit green colour in *E. coli* (Figure S2). Thus, we aimed to test whether the constructs were expressed at sufficiently high levels and also bound FMN.

### The iLOV^L470T/Q489K^ and iLOV^V392K/F410V/A426S^ mutants are expressed at low levels likely due to misfolding

To better understand the reasons behind the absence of fluorescence, we over expressed the proteins in *E. coli* and purified them. We then ran them on an SDS gel (Figure 3). While the WT and the iLOV^V392K^ mutant ran as single bands at around 15 kDa, higher molecular weight bands (~70–80 kDa) were observed for the iLOV^L470T/Q489K^ and iLOV^V392K/F410V/A426S^ mutants (Figure 3). Moreover, the iLOV^L470T/Q489K^ mutant had very low concentration.

**Figure 3:**
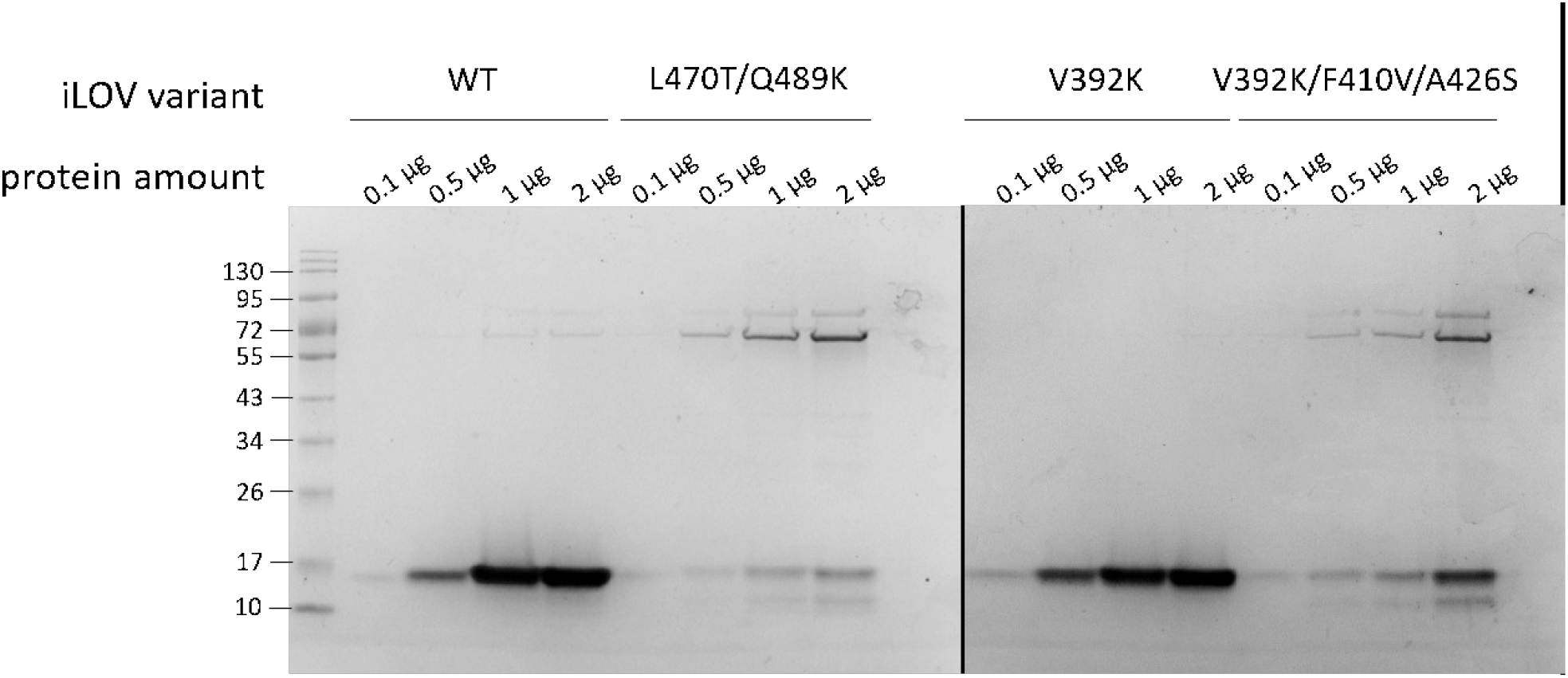
SDS gel of purified proteins. The indicated amounts of proteins were diluted in 10 μl storage buffer, boiled and run on a 10% SDS gel. The expected size of all proteins is 15.1 kDa. Pictures of gels were taken with a BIO-RAD ChemiDoc XRS+.

To investigate whether the higher molecular weight (MW) bands observed in the SDS gels for the double and triple mutants were constituted by the iLOV mutant themselves or were other proteins, which co-purified with the mutants, we ran mass spectrometry analysis (MS) of these bands. The results indicated that the high MW bands corresponded to the chaperones GroEL (57.34 kDa) and DnaK (also known as Hsp70; 69.13 kDa) running together with the iLOV mutants (15.1 kDa) leading to a total weight of 72.44 and 84.23 kDa, respectively. GroEL and DnaK have been previously detected in other studies with purified proteins from *E. coli*^23^. Being chaperones, they might be associated with newly synthesized proteins to help them fold properly. Interestingly, visual inspection of the purified proteins showed that only the WT protein fluoresced (Figure S3). To further clarify this observation, we conducted optical spectroscopy with the purified proteins.

### Spectroscopic analyses indicate that the V392K mutation leads to loss of FMN binding and, consequently, fluorescence

Next, we measured the absorption spectrum of all purified iLOV proteins between 250–800 nm. Surprisingly, the mutants bearing the V392K mutation, namely iLOV^V392K^ and iLOV^V392K/F410V/A426S^, did not show the peak at 450 nm typical of FMN. Considering the iLOV^V392K^ mutant was purified as single band as seen in the SDS gel (Figure 2), these results suggested that residue V392 is crucial for either the correct folding of the protein and/or its binding to FMN, which is essential for fluorescence. This was also in line with our cell and protein pellet observations (Figures S2 and S3). As no FMN absorption is observed for these mutants, we went on with the excitation-emission spectra measurements only for the iLOV^L470T/Q489K^ double mutant alongside the WT. In contrast to the QM/MM predictions of Khrenova and colleagues^20^, both excitation and emission spectra of iLOV^L470T/Q489K^ were slightly blue-shifted by ~2 nm (Figure 4). Next, quantum yields and the brightness of iLOV^L470T/Q489K^ were determined.

**Figure 4:**
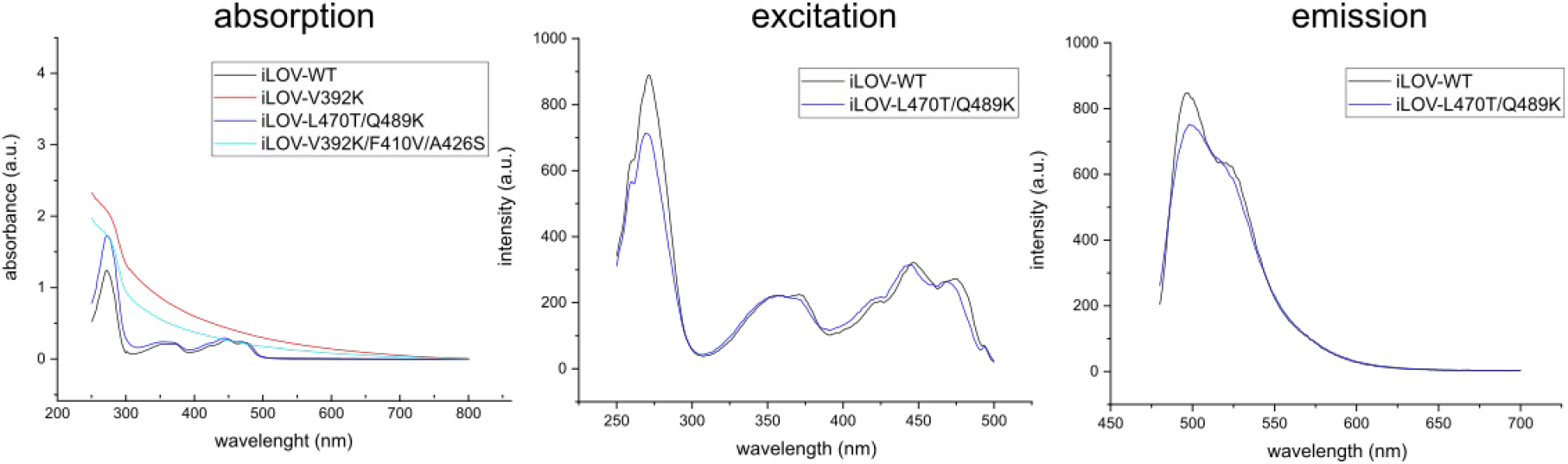
Absorption, excitation and emission spectra measurements of iLOV mutants. For all purified proteins, absorption spectra were measured between 250–800 nm. The excitation and emission spectra measurements were made between 250–500 nm and 480–700 nm, respectively.

Finally, we determined the quantum yield and the brightness of the iLOV^L470T/Q489K^ mutant and WT iLOV. The results for WT iLOV were in line with previous reports^11^. Interestingly, despite the low expression levels and the chaperone co-purification (Figures 2 and 3), iLOV^L470T/Q489K^ exhibited higher quantum yield and brightness compared to WT iLOV (Table 1).

**Table 1:** Quantum yield and brightness measurements of WT iLOV and iLOV^L470T/Q489K^. Numbers in parentheses are calculated by using the literature quantum yield of WT iLOV. (Linear regression plots used to calculate quantum yields are given in Figure S4).

|  | WT | iLOV<sup>L470T/Q489K</sup> |
| --- | --- | --- |
| **Quantum Yield ($\Phi_s$)** | 0.39 ± 0.06 (Literature 0.34<sup>11</sup>) | 0.47 ± 0.05 (0.40 ± 0.05) |
| **Brightness** | 4875 ± 750 (4250) | 5875 ± 625 (5000 ± 250) |

To further mechanistically explore the discrepancy between the computational predictions of Khrenova and colleagues^20^ and our experimental results, we used the DUET tool^24^, reported to be the best ranking tool in recent benchmark analysis^25^ to investigate the stability effect of single point mutations on the protein structure. Confirming our chaperone co-purification and lack of FMN binding observations, the double (iLOV^L470T/Q489K^) and triple (iLOV^V392K/F410V/A426S^) mutants were predicted to become unstable upon introduction of the mutations (see Table 2). Additionally, we used PremPLI^26^, a protein structure-based machine learning method for the prediction of the effects of single point mutations on ligand binding affinity. We analysed the change of FMN binding affinity to the iLOV protein upon introduction of the mutations used in our study. In line with our observation of lack of FMN binding for constructs bearing the V392K mutation, PremPLI^26^ indicates the strongest decrease (1.05 kcal mol^-1^) in the binding affinity towards FMN for mutants with this mutation (Table 2). Taken together, our results suggest that applying simple *in silico* stability and ligand binding analyses could help determine mutations that would likely lead to fluorescence loss.

**Table 2:**
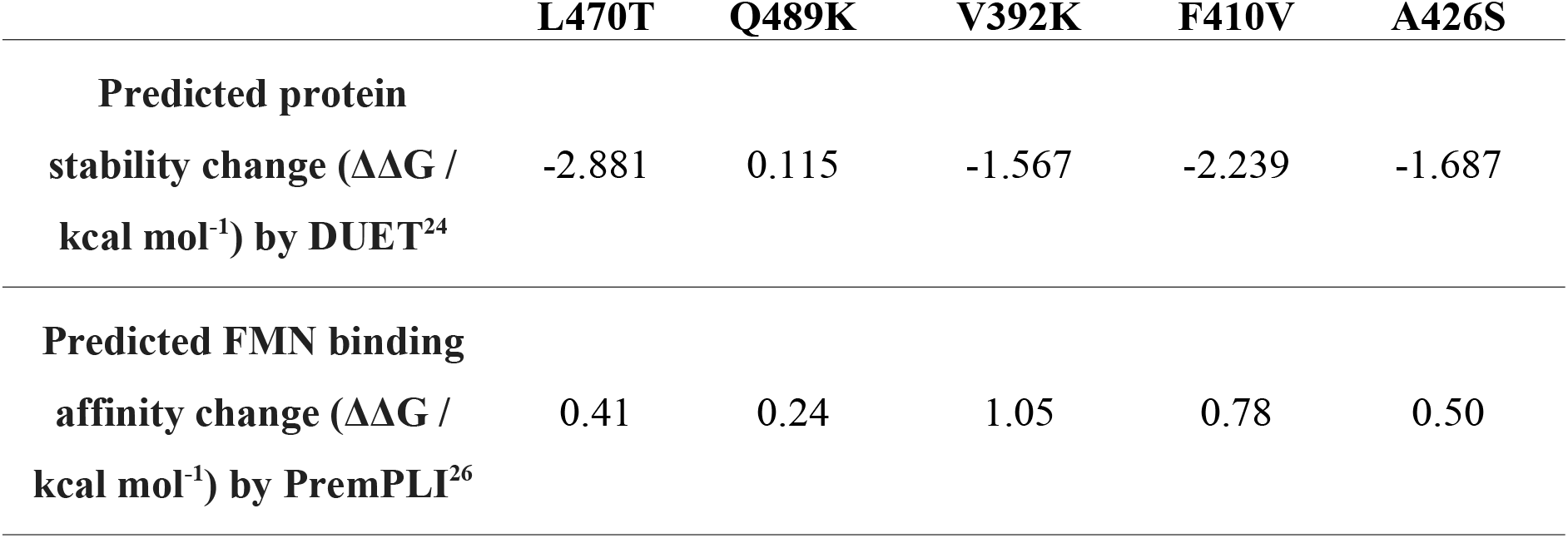
*In silico* analyses of iLOV structure. Protein stability change predictions of DUET^24^ and FMN binding affinity change predictions of PremPLI^26^ by the introduction of single point mutations to WT iLOV structure

## Discussion

In this study, we set out to analyse the excitation and emission spectra of two iLOV mutants, iLOV^L470T/Q489K^ and iLOV^V392K/F410V/A426S^, which were computationally predicted to be red-shifted of about 50 nm^20^. We found that the iLOV^V392K/F410V/A426S^ mutant does not bind FMN and that the iLOV^L470T/Q489K^ mutant is slightly blue-shifted in both excitation and emission spectra.

Initially, Khrenova and colleagues predicted the single mutant iLOV^Q489K^ to be shifted in excitation and emission maxima by 52 and 97 nm, respectively^18^; however, when experimentally measured, the excitation and emission maxima exhibited rather a blue-shift of about 10 nm^19^. This highlighted a disagreement between the predictions and the experimental validations. Even though the QM/MM modeling approach was subsequently updated^20^, our results argue that it is still not able to completely capture the complexity of the interaction between the iLOV^L470T/Q489K^ mutant and its chromophore. Nevertheless, the iLOV^L470T/Q489K^ mutant has ~20% increased brightness and quantum yield compared to WT iLOV.

Several studies recently also provided a molecular characterization of the spectral effects of mutations on iLOV to gain a mechanistic understanding^27–29^. Röllen et al.^29^ have shown that iLOV^V392T/Q489K^ fluorescence has a slight red-shift with an emission spectrum maximum of 502 nm. Its homolog mutant CagFbFP^I52T/Q148K^ (emission maximum 504 nm) and another blue-shifted variant CagFbFP^Q148K^ (emission maximum 491 nm) were used in fluorescence microscopy experiments and were successfully spectrally separated. While iLOV^V392T/Q489K^ (and its homolog CagFbFP^I52T/Q148K^) was mutated at the same sites as in our study, the V392T mutation does not reduce the affinity of iLOV for FMN as much as V392K does (FMN ΔΔG for V392T 0.67 kcal mol^-1^ vs for V392K 1.05 kcal mol^-1^ predicted by PremPLI^26^). For the rational design of iLOV variants, using *in silico* predictors could guide researchers to filter out mutations that would disrupt protein stability and/or reduce FMN binding.

Previous efforts to improve the properties of iLOV for imaging purposes resulted in the generation of phiLOV, an iLOV derivative which possesses a superior photostability, thus tackling one of the major drawbacks of WT iLOV^30^. Furthermore, iLOV variants with improved brightness recently have been reported^14,15^. It remains to be seen whether future red shift mutagenesis efforts of iLOV can be combined with these more stable and brighter versions of iLOV.

## Material and Methods

### Plasmid construction

Constructs were generated using classical restriction enzyme cloning or golden gate cloning. The *ilov* gene was amplified from iLOV-N1, gift from Michael Davidson (Addgene plasmid #54673; http://n2t.net/addgene:54673; RRID:Addgene_54673) and cloned into pcDNA3.1(+) using HindIII and EcoRI restriction sites, yielding plasmid PL01. PL01 was used as a template to clone all iLOV variants used in this study. Single-point mutations Q489K and V392K were created via overhang PCR yielding PL02 and PL04, respectively. The L470T mutation was introduced into PL02 using golden gate cloning. Amplicons were digested with HindIII and BpiI, or BpiI and EcoRI, respectively, and cloned into pcDNA3.1(+) yielding PL03. The A426S mutation was inserted into PL04 by addition of a XbaI restriction site into the coding sequence and cloned into pcDNA3.1(+) yielding PL05. The F410V mutation was introduced using golden gate cloning. Amplicons were digested with HindIII and BpiI or BpiI and EcoRI, respectively, and cloned into pcDNA3.1(+) yielding PL06.

For expression in and purification from *Escherichia coli*, the *ilov, ilov^L470T/Q489K^*, *ilov^V392K^* and *ilov^V392K/F410V/A426S^* genes were amplified and then cloned into pET28a using NdeI and XhoI restriction sites, yielding PL07-PL10. The sequences of all constructs were verified using Sanger sequencing.

### Bacterial cell culture and transformation

Bacterial strain used in this study was *E. coli* Rosetta (DE3) pLysS. iLOV constructs were transformed into chemically competent cells via heat shock (42 °C, 1.5 min), plated on lysogeny broth (LB) agar plates containing 0.05 mg/mL kanamycin and incubated overnight at 37 °C. For expression, 5 mL liquid LB were inoculated with a single colony from a fresh LB agar plate and incubated overnight at 37 °C and 200 rpm. The next day, the cultures were used to inoculate 1 L of LB with a starting OD_600_ of 0.1. All liquid media were supplied with 0.05 mg/mL kanamycin.

### Protein expression and purification

Each pET28a plasmid harbouring one iLOV construct was transformed into *E. coli* Rosetta (DE3) pLysS cells and cultures were grown at 37 °C and 220 rpm until they reached an OD_600_ of 0.4. IPTG (Isopropyl-β-D-thiogalactopyranosid) was then added to the culture to a final concentration of 1 mM, and the culture was further grown at 18 °C for 16 h. Afterwards, cells were harvested via centrifugation at 5000 × g for 30 minutes. The pellets were resuspended in lysis buffer (50 mM KH_2_PO_4_ pH 7.5, 300 mM NaCl and 10 mM imidazole pH 8.0) containing one tablet of cOmplete protease inhibitor cocktail (Roche), and lysed by sonication. The lysate was clarified by centrifugation at 20,000 x g for 30 minutes at 4 °C and loaded onto an IMAC nickel column (1 mL) using the Bio-Rad NGC automated liquid chromatography system. The column was washed with a wash buffer (same as lysis buffer but with 20 mM imidazole and 10 % glycerol) and eluted with elution buffer (same as lysis buffer with further addition of 10 % glycerol and 500 mM imidazole). Finally, the elution buffer was replaced with a storage buffer (50 mM HEPES-KOH pH 7.25, 150 mM KCl, 10 % glycerol and 0.1 mM EDTA pH 8.0) using a P-6 desalting column (10 mL).

### Fluorescence microscopy

Fluorescence microscopy was performed on a Zeiss Axio Observer Z1/7 (Carl Zeiss, Germany) inverted wide-field microscope, equipped with a Colibri 7 LED light source, an Alpha Plan-Apochromat x 100/1.46 oil DIC (UV) M27 objective, filter sets 38 HE (ex. 450-490, dichroic beamsplitter495, em. 500-550; sfGFP) and an Axiocam 506 Mono camera. 6 μL of culture (OD_600_ = 1.5) were loaded on a glass slide.

### SDS gel and mass spectrometry analyses of purified iLOV proteins

The concentration of purified proteins was determined using NanoDrop One (Thermo Scientific). For performing SDS-PAGE, protein samples were set to contain 0.1 μg, 0.5 μg, 1 μg and 2 μg, mixed with Laemmli buffer and incubated at 95 °C for 15 minutes. After incubation, the samples were spun down and 20 μL were loaded on a pre-cast 12 % agarose gel (MiniProtean TGX gel, BIO-RAD). SDS-PAGE was performed at 130 V for 40 minutes. Afterwards, gels were stained for 1 h using ReadyBlue^TM^ Protein Gel Stain (Sigma-Aldrich) and destained with ddH_2_O overnight. Images were taken with the ChemiDoc^TM^ XRS+ system (BIO-RAD). Protein bands analysed using mass spectrometry were excised, treated with 10 mM dithiothreitol and 10 mM iodoacetamide and subsequently digested with trypsin. The peptide solution was then separated on a Waters Acquity I-class UPLC in positive HD-MSE mode using a Waters Peptide CSH C18 column (2.1 mm x 150 mm, 1.7 μm particle size). A gradient from 1% to 40% ACN/0.1% formic acid (v/v) in water/0.1% formic acid (v/v) was utilized at a starting temperature of 80 °C and dissolving temperature of 400 °C with a gas flow rate of 800 l/h. Spectra obtained by the separation were analysed by matching with the UniProt database.

### Spectroscopy analyses

All optical spectra were measured at room temperature in a fluorescence cuvette with 1 cm path length (Art. No. 105-250-15-40, Hellma Analytics). All samples (volume 100 μL) with a concentration of 500 μg/mL were dissolved in a storage buffer (50 mM HEPES-KOH, pH 7.25, 150 mM KCl, 10 % (v/v) glycerol, 0.1 mM EDTA). Absorption spectra were recorded from 250–800 nm (UV-2450 Spectrophotometer, Shimadzu). Fluorescence measurements were performed with a Luminescence Spectrometer (LS 55, Perkin Elmer). Fluorescence excitation spectra were recorded from 250–500 nm at a fixed emission wavelength of 496 nm. Emission spectra were measured from 480–700 nm at an excitation wavelength of 450 nm. The quantum yields were determined with the comparative method^31,32^ using FMN dissolved in a storage buffer as reference. For the linear regression the integrated emission (480–700 nm) was plotted against the absorbance at 450 nm. The quantum yield was then calculated using the following equation:

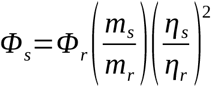

where *m* indicates the slope from the linear regression and *η* the refractive index of the sample or the reference, respectively. The fraction of refractive indices equals to 1 since the same buffer was used. For the quantum yield Φ*r* of FMN 0.24 was used^33^. The brightness was calculated as the product of the determined quantum yield and the extinction coefficient of free FMN as 12500 M^-1^ cm^-1^^34^.

### In silico protein stability and FMN binding affinity analyses of iLOV mutants

WT iLOV (for amino acid sequence see Supplementary Information) was modeled by the Swiss-Model web-server^21^ using PDB id: 4EEP as template. The resulting structure was submitted to DUET^24^ and PremPLI^26^ web-servers to predict the change of protein stability and FMN binding affinity, respectively.

## Acknowledgments

We thank Lena Appel and Matthias Boll (University of Freiburg, Germany) for their mass spectrometry analyses and Alexander Löwer (Max Dellbrück Center for Molecular Medicine, Germany) for sharing the H1299 cell line, and Michael Davidson (Florida State University, USA) for sharing iLOV-N1 vector with us.

## Supplementary Information

### ilOV variants

Amino acid sequences of the iLOV variants used in this study are given below:

**iLOV(WT):**MFIEKNFVITDPRLPDNPIIFASDGFLELTEYSREEILGRNARFLQGPETDQATVQKIRDAIRDQRETTVQLINYTKSGKKFWNLLHLQPVRDQKGELQYFIGVQLDGS DHV
**iLOV^L470T/Q489K^**: MFIEKNFVITDPRLPDNPIIFASDGFLELTEYSREEILGRNARFLQGPETDQATVQ KIRDAIRDQRETTVQLINYTKSGKKFWNL**T**HLQPVRDQKGELQYFIGV**K**LDGSDHV
**iLOV^V392K/F410V/A426S^**: MFIEKNF**K**ITDPRLPDNPIIFASDG**V**LELTEYSREEILGRN**S**RFLQGPETDQATVQ KIRDAIRDQRETTVQLINYTKSGKKFWNLLHLQPVRDQKGELQYFIGVQLDGSDHV
**iLOV^V392K^**: MFIEKNF**K**ITDPRLPDNPIIFASDGFLELTEYSREEILGRNARFLQGPETDQA TVQKIRDAIRDQRETTVQLINYTKSGKKFWNLLHLQPVRDQKGELQYFIGVQLDGSD HV

### Mammalian cell culture and transient transfection

Human non-small cell lung carcinoma cells (H1299; kindly provided by Alexander Loewer, Max Delbrück Center for Molecular Medicine, Berlin) were maintained in RPMI 1640 media supplemented with 10% foetal bovine serum, 100 U/ml penicillin and 100 μg/ml streptomycin. Cells were cultivated at 37 °C and 5% CO_2_ and were passaged when reaching ~90% confluence.

For microscopy experiments, H1299 cells were seeded in μ-Slide 8-well chambers (Ibidi GmbH) at densities of ~15000 per well. The following day, the samples were transfected with 150 ng of construct DNA using Lipofectamin 2000 (ThermoFisher Scientific) according to the manufacturer’s manual. Samples were imaged 36 hours post-transfection.

### Fluorescence microscopy of mammalian cells

Mammalian cells were imaged at 37 °C and 5% CO_2_ in μ-Slide 8 well chambered coverslips (Ibidi GmbH) on a Zeiss AxioObserver wide-field microscope equipped with an incubation chamber, a 40x air objective (0.7 NA), the Colibri.2 LED light source and a filter set for eGFP (Figure S1).

**Figure S1:**
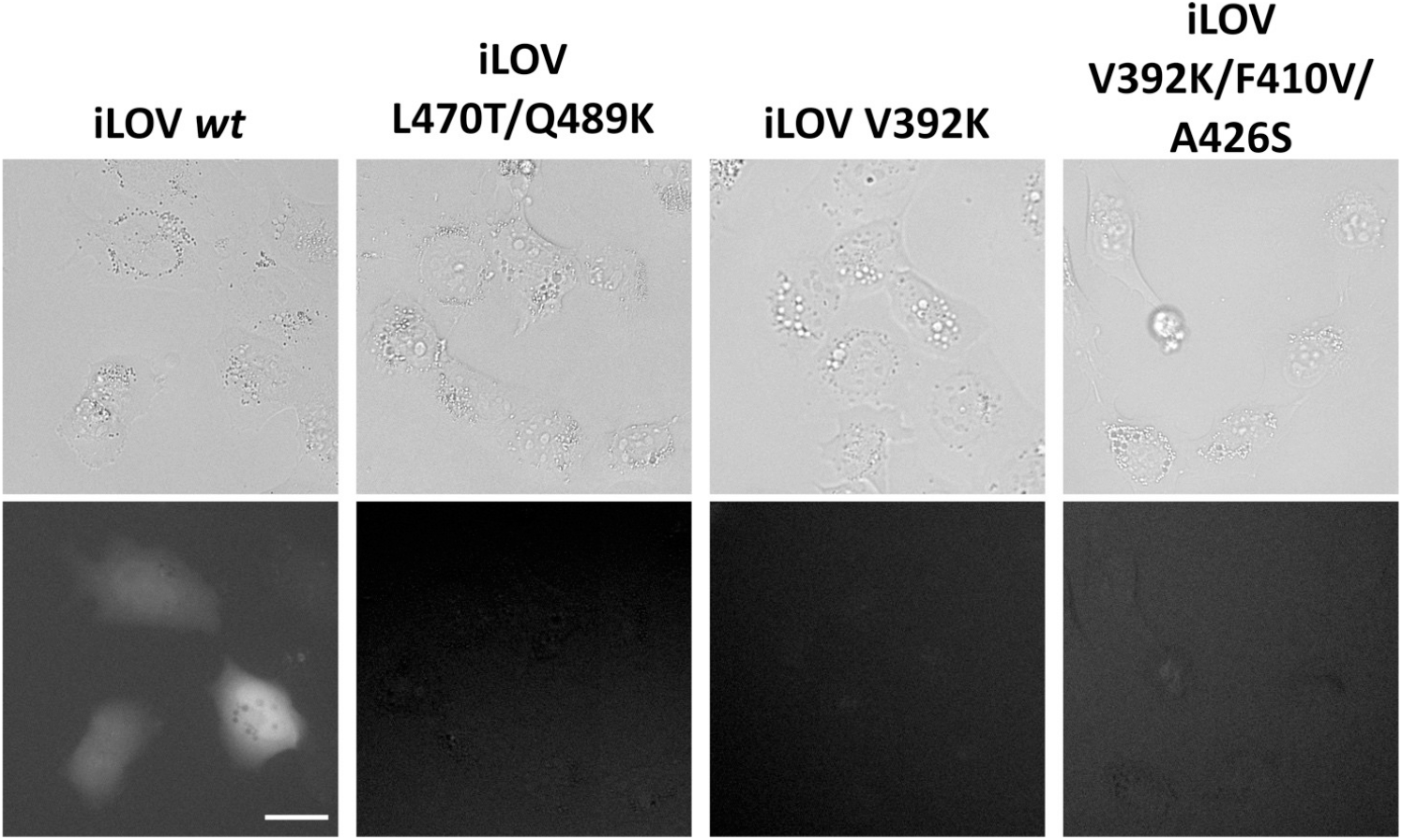
Analysis of iLOV variants in fluorescence microscopy. Representative bright field (top) and eGFP (bottom) images of H1299 cells transiently transfected with the indicated iLOV construct. Scale bar for all micrographs is 30μm.

**Figure S2:**
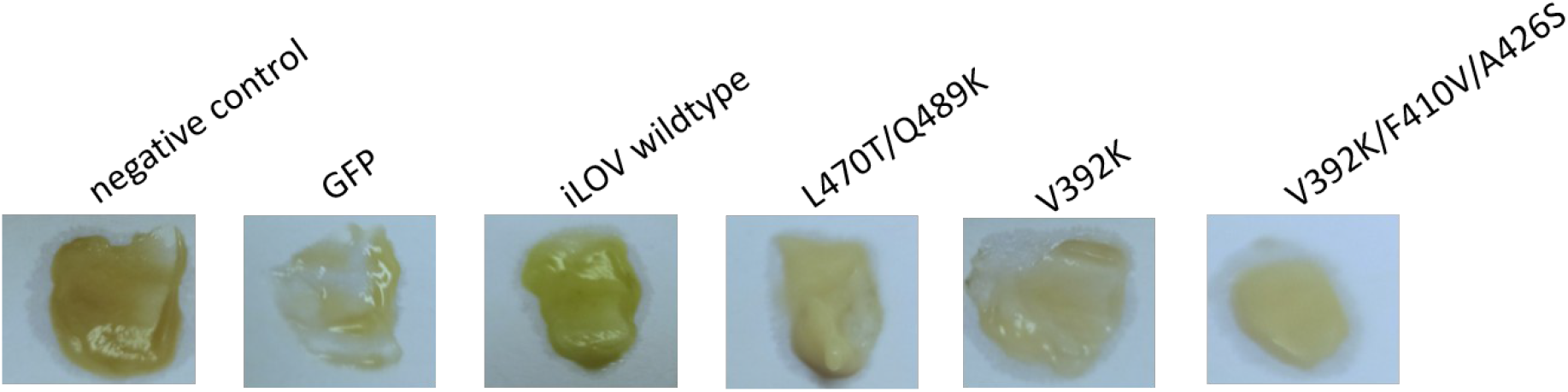
Comparison of the pellets of iLOV expressing cells. Representative picture of E. coli Rosetta^TM^(DE3) pLysS cells transfected with the indicated iLOV construct. Negative control is non-transfected cells. Cells transfected with a plasmid expressing GFP serve as a positive control. A 100 mL culture was grown to saturation while expressing iLOV variants and then harvested at 5000 x g for 20 minutes. 20 mg of the formed pellet was streaked on white paper.

**Figure S3:**
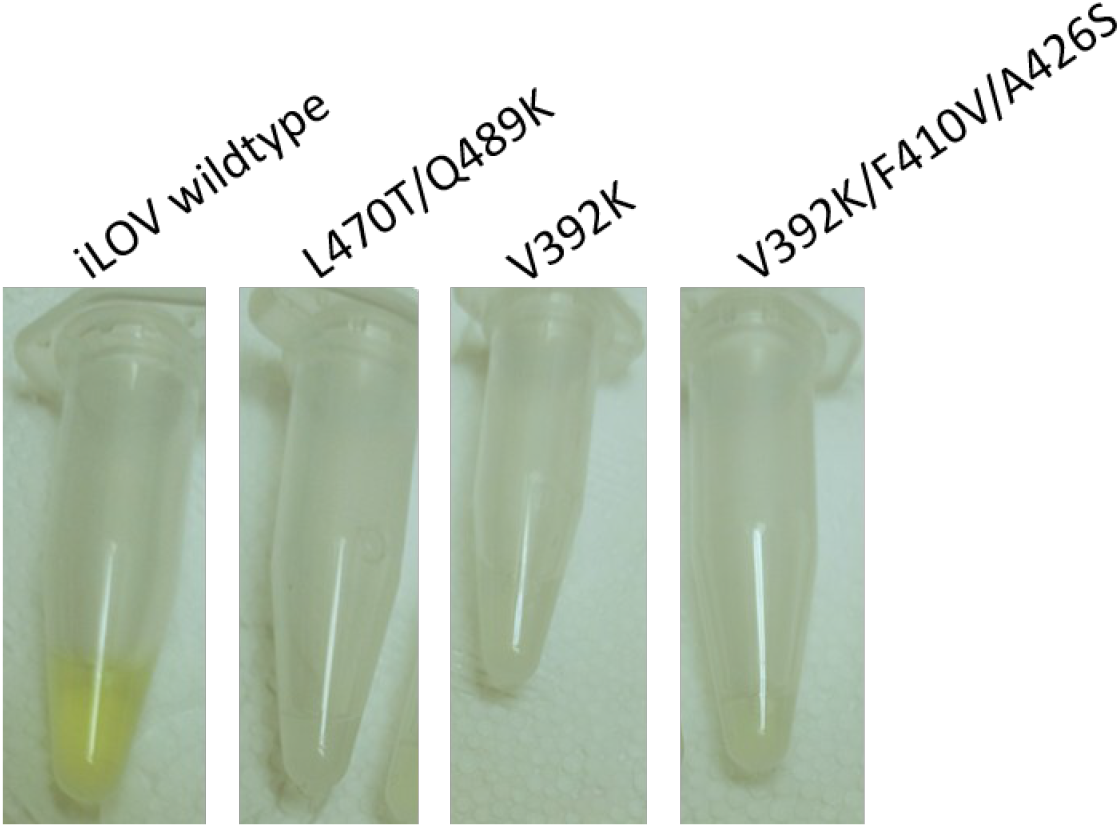
Comparison of the colour of iLOV-variants. Representative picture of purified iLOV proteins. Samples are set to contain 1 mg/ml of proteins in an 1.5 mL Eppendorf tube.

**Figure S4:**
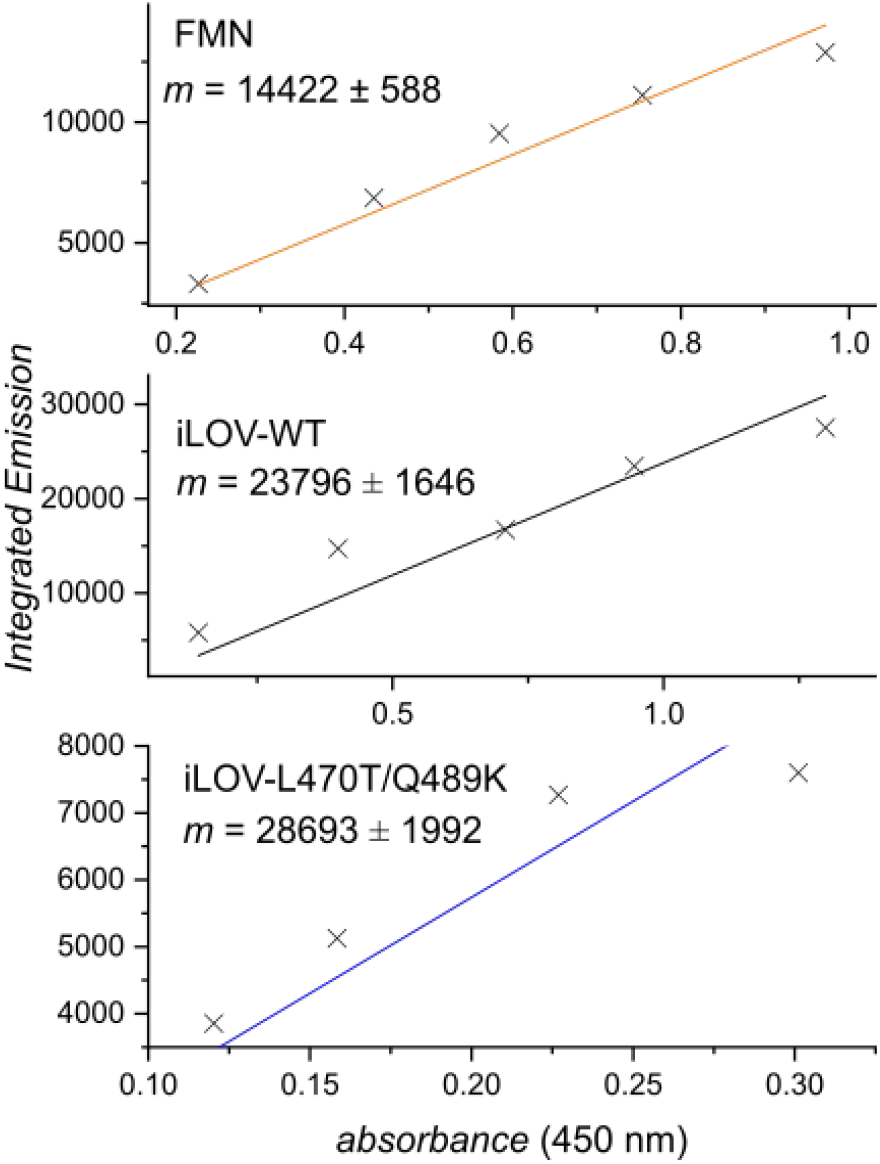
Comparative linear regression analysis. The integrated emission (480–700 nm) was plotted against the absorbance at 450 nm to calculate quantum yields given in Table 1.

